# Interpreting Inverse Correlation Time: from Blood flow to Vascular Network

**DOI:** 10.1101/2022.07.15.500238

**Authors:** Qingwei Fang, Chakameh Z. Jafari, Shaun Engelmann, Alankrit Tomar, Andrew K. Dunn

## Abstract

The inverse correlation time (ICT) is a key quantity in laser speckle contrast imaging (LSCI) measurements. Traditionally, ICT is regarded as a metric of blood flow, such as speed or perfusion. However, we highlight that ICT not only contains important information about blood flow, but also reflects the underlying structure of the vascular network. In the past, ICT has been found to be correlated with vessel diameter. Here, we further report that ICT exhibits a different sensitivity to blood flow depending on vessel orientation. Specifically, ICT is more sensitive to blood flow speed changes in vessels descending from or arising to the tissue surface, compared with those laying parallel to the surface. Those findings shift our understanding of ICT from purely blood flow to a combination of blood flow and vascular network structure. We also develop theories to facilitate the study of vascular network’s impact on ICT.

## 1. Introduction

Laser speckle contrast imaging (LSCI) has gained attention in recent years for the quantification of blood flow in biomedical imaging applications^1–3^. It is non-invasive, non-ionizing and label free. In general, the higher the blood flow speed, the more rapidly the speckle patterns will vary in time, leading to a lower speckle contrast when integrated over the camera exposure time^4,5^. Valuable information about blood flow perfusion in regions of interest can be extracted in real time from the continuous and wide-field 2-dimensional (2D) monitoring of speckle contrast^6,7^. Nevertheless, the speckle contrast only provides qualitative measurements of the underlying blood flow^8^. Numerous efforts have been put into advancing LSCI from qualitative to a quantitative imaging modality^9–15^.

One promising path is to relate speckle contrast to the autocorrelation function of detected electric field, *g*_1_(*τ*) and extract the inverse correlation time (ICT) as the index of blood flow. Within a limited set of conditions, ICT is proportional to the typical speed of blood flow within a certain range^16,17^, and it has demonstrated great potential in quantifying cerebral blood flow and facilitating intraoperative flow monitoring^18–21^.

In view of the promise of ICT in transforming LSCI to a quantitative imaging modality, the strategy to extract ICT from measured speckle dynamics is gaining the significant attention. Initially this was done with single-exposure LSCI^16,17^, and more recently, multi-exposure speckle imaging (MESI) was developed to extract ICT more accurately from the confounding effects of static scattering, instrumentation noise and loss of correlation due to speckle averaging^22–24^. Recently, dynamic light scattering imaging (DLSI) was proposed to reduce the ICT estimating error owing to an inaccurate model of electric field autocorrelation function^25^.

Though ICT has been mainly interpreted as a metric of blood flow, such as speed, or perfusion, there is increasing evidence that ICT is subject to the structure of the vascular network. Kazmi et al.’s experimental results showed that ICT might be quantifying neither volumetric flux nor flow speed but the product of the flow speed and vessel diameter^26^. Fredriksson et al. reported the vessel packaging effect in which the confinement of blood to vessels with an average diameter of 40 *μ*m could lead to a 50% reduction in perfusion estimation by LSCI compared with homogeneous blood distribution inside the tissue^27^. Jafari et al. found that ICT could be shifted over 10 times under the homogeneous assumption compared with using the actual vascular geometry, highlighting the significant impact of vascular geometry on ICT^28^.

The impact of vessel orientation on ICT has not been fully investigated yet. In this work, we conducted Monte Carlo simulations and found that ICT of vessels perpendicular to the tissue surface, i.e. descending from/arising to the surface, exhibits a higher sensitivity to blood flow changes than that of vessels laying on the surface. Such finding is confirmed by experimental validation *in vivo* combining MESI and 2-Photon (2P) imaging. Those results suggest that the structure of the underlying vascular network deserves more attention than it currently receives in the interpretation of ICT. We also develop a generalized theory to facilitate the study on the impact of vascular network structure. It is compatible with all extant g_1_(τ) models and free of assumptions about groundtruth blood flow speeds.

## 2. Theory and Methods

### 2.1. A unified theoretical framework for ICT interpretation

According to the traditional dynamic light scatteirng (DLS) and diffusing wave spectroscopy (DWS) theories, in different scattering regimes (single vs. multiple) and flow conditions (ordered vs. unordered), different assumptions should be made about the form of the electric field autocorrelation function *g*_1_(*τ*) [29–33], as shown in **Fig. 1A** [25]. *τ_c_* is the correlation time and ICT is defined as 1/*τ_c_*. The modulation number *n* is the main differentiating factor among those models.

**Figure 1:**
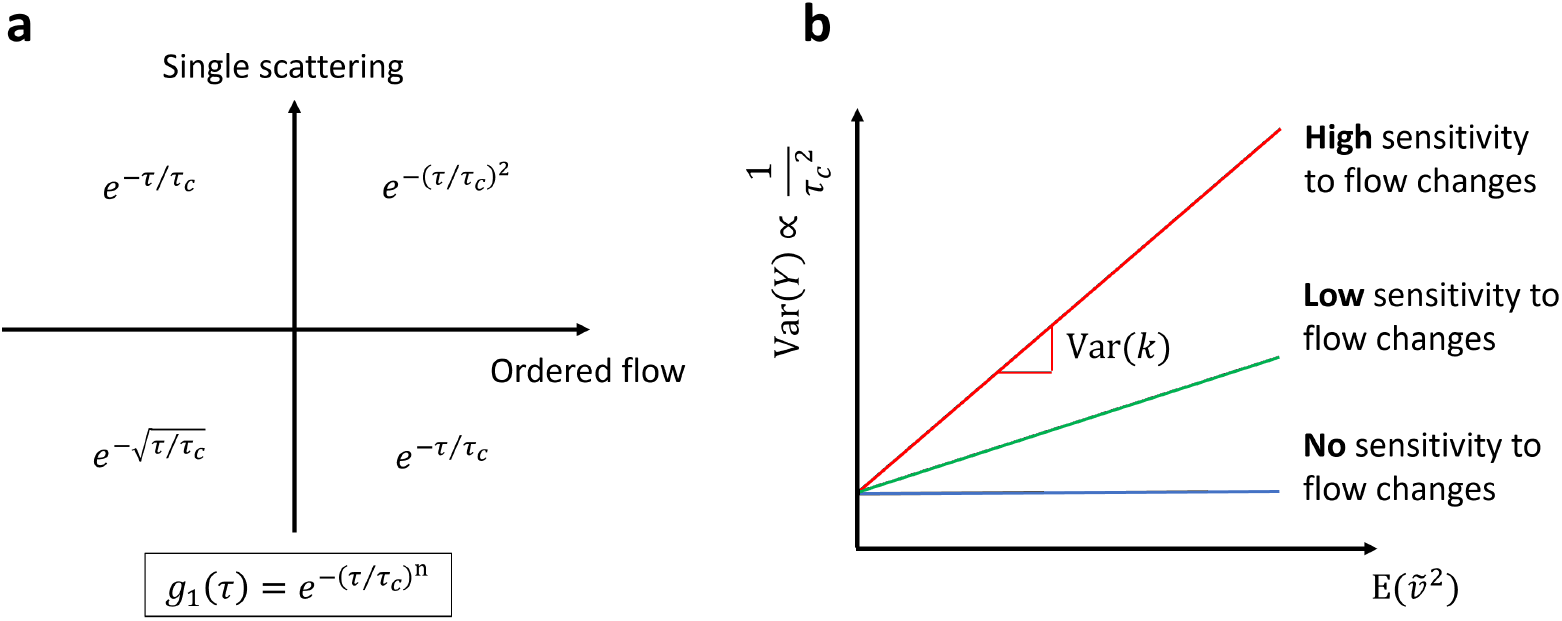
A unified theoretical framework for ICT interpretation. **a** The various *g*_1_(*τ*) models proposed in treatment of the complex vascular network in LSCI. Extant models of the electric field auto-correlation function in various cases of scattering regimes and flow conditions. *τ_c_* is the correlation time and ICT is defined as 1/*τ_c_*. On the quadrant plot, the opposite of the order motion is unordered motion (also known as diffusive motion) and that of single scattering is multiple scattering. **b** Physical meaning of the characteristic variance of dynamic scattering. The vertical axis depicts the variance of dynamic scattering Var (*Y*) which is proportional to the square of ICT. The horizontal axis is the weighted average of blood flow speeds sampled by the photon scattering process. The slope of the plot is Var(*k*). A steeper slope indicates a vascular network of higher sensitivity to flow changes.

In this paper, we first introduce a unified theoretical framework that is compatible with all extant *g*_1_(*τ*) models. After Monte Carlo simulation of photon migration inside the tissue, the electric field auto-correlation function *g*_1_(*τ*) can be calculated according to Eq. 1 (ref. [34])

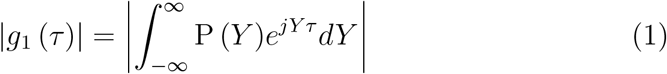

where 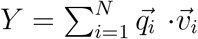 and 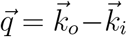. *N* is the number of dynamic scattering events experienced by a single photon. N can be larger than 1 hence Eq. 1 accommodates both single and multiple scatter regimes. 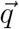 is the momentum transfer vector defined as the difference between the scattering wave vector 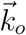 and the incident wave vector 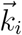 [35]. The direction of 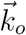 and 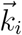 is the propagating direction of the scattered light and incident light, respectively. The amplitudes are both equal to *k*_0_, the wavenumber. Finally, 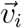 is velocity of the scattering particle in the *i*-th dynamic scattering event experienced by the photon. For clarity sake, speed refers to scalars while velocity corresponds to vectors in this paper. Note that since the instantaneous velocity of Brownian particles is physically measurable^36^, we expand the definition of 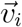 from the velocity of ordered motion as defined in ref. [34] to include both ordered and diffusive motions. *P*(*Y*) is the probability density function of *Y*. When some photons generate the same *Y* value, their weights will be added together to give rise to *P*(*Y*). *Y* can be interpreted as the accumulation of frequency shifts induced by dynamic scattering (see Supplemental Material section 1 for a more concise proof than ref. [37]).

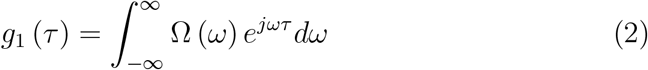

With *g*_1_(*τ*) rewritten in terms of *Y*, we notice that *g*_1_(*τ*) is the Fourier transform of *P*(*Y*). Considering the Wiener-Khintchine theorem, if Ω (*ω*) is the normalized power spectral density of the detected electric field, then *g*_1_(*τ*) would be the Fourier transform of Ω (*ω*) (Eq. 2) (ref. [38]). Therefore, we arrive at Eq. 3 that *P*(*Y*) is a shifted version of Ω (*ω*)

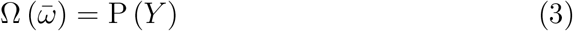

where 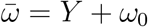 and *ω*_0_ is the center frequency of the electric field.

Substituting *P*(*Y*) for Ω(*ω*) is advantageous in that *P*(*Y*) reveals the mechanism of spectrum broadening in LSCI since *Y* is the accumulation of frequency shifts induced by dynamic scattering. *P*(*Y*) and specific forms of *g*_1_(*τ*) can be bridged by Eq. 4 and 5. For *g*(*ρ*) = *e*^−|*τ*|/*τ_c_*^,

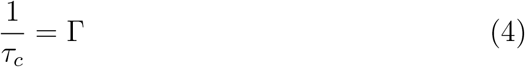

where Γ is the half width at half maximum of the Lorentzian 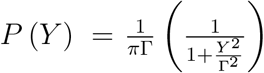. For *g*_1_(*τ*) = *e*^−(*τ*/*τ_c_*)2^,

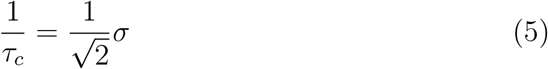

where *σ* is the standard deviation of the Gaussian 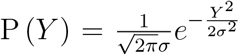. See proof in Supplemental Material section 2.

These two equations (Eq. 4 and 5) provide new interpretations of ICT (i.e., 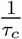) from the point of view of *P*(*Y*). For Lorentzian *P*(*Y*), ICT determines the half width at half maximum while in Gaussian *P*(*Y*), ICT is a scaled version of the standard deviation. Note that in both cases, ICT serves as a kind of measure of the broadening of the spectrum of *Y*. In fact, as long as the spectrum of detected light is both bandwidth- and amplitude-limited, a more general linear relationship

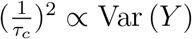

can be shown to hold for any electric field auto-correlation functions taking the form of *g*_1_(*τ*) = *e*^−(|*τ*|/*τ_c_*)*n*^ (see Supplemental Material section 3 for proof). This suggests that ICT in the various *g*_1_(*τ*) models shown in Fig. 1a can be unified as the reduced case of the standard deviation of *Y*. Since *P*(*Y*) is essentially the same as the power spectral density of the electric field of detected light (Eq. 3), ICT can be interpreted as the scaling factor of the variance of the detected optical spectrum as well. Since Var (*Y*) measures the variance of frequency shifts in the dynamics scattering process, we call it the variance of dynamic scattering.

### 2.2. Characterizing the inherent properties of vascular network to produce ICT

In this section, we aim to decouple the effects of blood flow speeds and vascular structure on ICT and develop tools to study the vascular network individually. Further analyzing the structure of *Y* and separate the contributions to *Y* by the blood flow and others, we arrive at Eq. 6

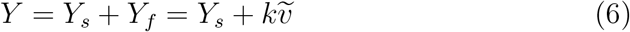

where *Y_f_* represents the contributions to *Y* by motions induced by blood flow while *Y_s_* denotes the component in *Y* that is independent of the blood flow. *Y_f_* can be expressed as the product of *k* and 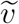 where 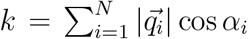, 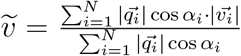 and *α_i_* is the angle between 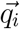 and 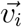. The 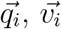 and *N* follow same definition as in Eq. 1.

Note that 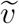 represents the weighted average of absolute blood flow speeds and the weighting is determined by the photon’s scattering geometry, i.e., the scattering angle and the angle between the momentum transfer vector 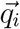 and velocity vector 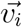. *k* is the sum of weights and equal to *Y_f_* when blood flow speeds are all unit speeds.

Also notice that *k* and 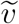 are well de-correlated from each other. To understand this, it is helpful to view *k* as the sum of weights and 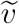 as the weighted average of blood flow speeds. When the sum of weights changes, we would not be able to predict how the weighted average will change if little is known about how the individual weight changes. The weighted average can increase or decrease. It can also remain invariant if the weights increase/decrease by the same ratio.

Furthermore, if we assume that blood flow speeds are uncorrelated with the photon’s scattering geometry (i.e., the scattering angle and the angle between 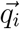 and 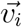 are uncorrelated with the magnitude of the flow vector), 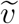 and *k* would be uncorrelated (see proof in Supplemental Material section 4). With this uncorrelated relationship, we arrive at Eq. 7 and 8

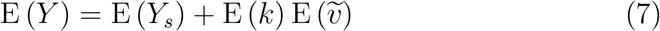

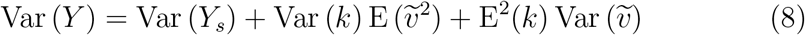

where E (*X*) and Var (*X*) give the expectation and variance of the random variable *X*, respectively.

Eq. 7 and 8 points out the relationship between statistical properties of *Y* and those of *k* and 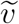. Further approximations to Eq. 8 can be made to accommodate blood flow quantification. Either of the following two conditions is sufficient for Eq. 9 to be applied with good accuracy. First, the vascular structure sampled by detected photons is dominated by a single vessel, i.e., 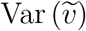 is approximately 0. This is true for major surface vessels where dynamic scattering events of detected photons are strongly localized within that vessel^8,21,33^. Second, the scattering of collected photons is fully randomized, i.e., E(*k*) is 0. This is true for parenchyma regions where dynamic scattering is dominated by micro-vessels whose orientation is randomized. Our simulation results of E (*k*) on parenchyma regions also support that as section 3.1 will show.

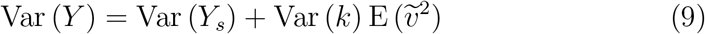

With Eq. 9, we define the characteristic variance of dynamic scattering as Var (*k*). If we plot Eq. 9 (Fig. 2), the physical implications of Var (*k*) are evident as the slope. Var (*k*) can be defined for each detection point (i.e., a pixel on the camera) in LSCI and it characterizes the ability of the sampled vascular network to decrease speckle contrast or increase ICT under the specified illumination/detection setup at the detection point. The larger the Var (*k*), the stronger the ability of the vascular network to decrease speckle contrast or increase ICT under the same blood flow speed, at the detection point. In addition, Var (*k*) is independent of the blood flow speed since it is defined with unit speeds. Hence, it reveals the inherent properties of the vascular network under the specified illumination/detection setup.

**Figure 2:**
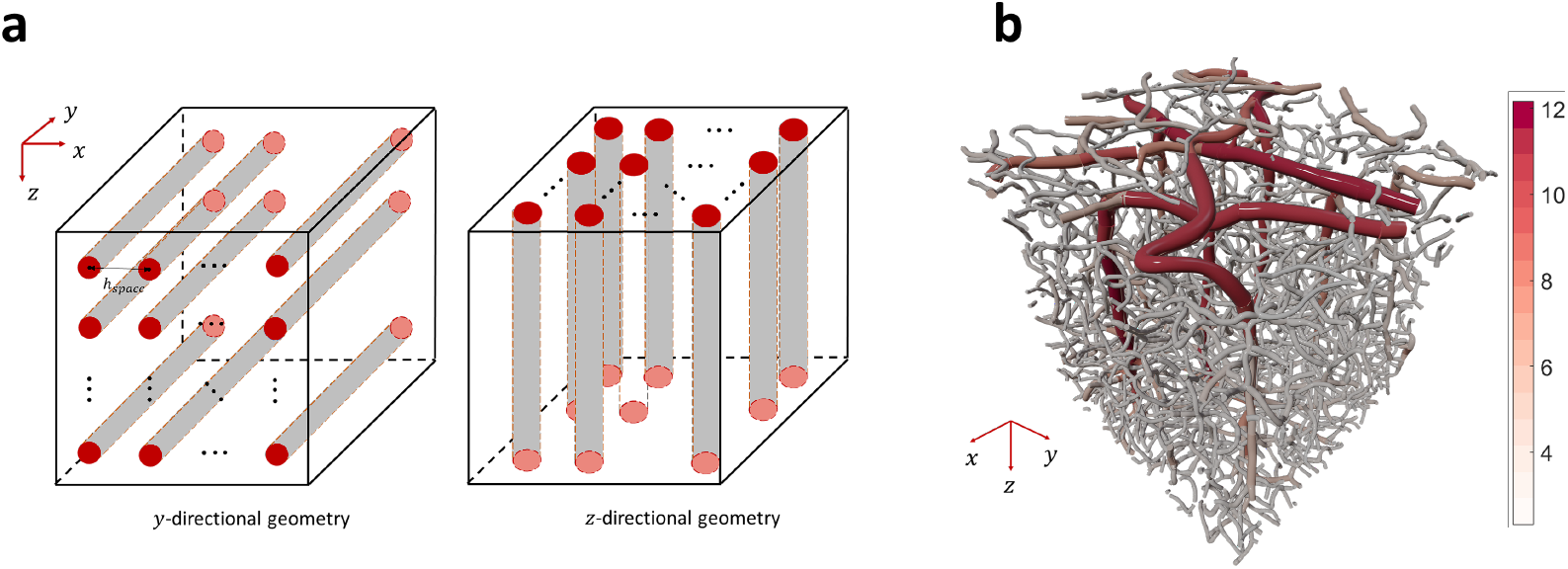
Vascular geometry in Monte Carlo simulation. **a** The diagram of simple *y*- and *z*-directional vascular geometry. **b** The realistic vascular geometry scanned from a mouse’s cerebral cortex. The geometry size is 554 × 554 × 606 *μ*m^3^. 3D rendering by Blender 2.81 (ref. [41]). Color coded by vessel radius (unit: *μ*m).

### 2.3. Monte Carlo simulation

The simulation algorithm is based on ref. [28]. The uniform flat beam profile is used for simulation. The effects of ordered motion of the RBCs along the direction of vessels are investigated in this study. Though the diffusive motion of RBCs has been recently suggested as dominating the correlation decay in diffuse correlation spectroscopy (DCS) measurements^39^, it is unclear whether the effects are due to the radially diffusive motion of RBCs within vessels or axially ordered motion of RBCs along the vessel but within a diffusive vascular network where vessels are curved and direction-randomized. Nor it is clear whether the observation holds in LSCI. Jafari et al. found that the axially ordered motion of RBCs along the vessel adequately describes the particle dynamics in LSCI given the strong agreement between experimental and simulated speckle contrast values^28^.Hence, the effects of radially diffusive motion of RBCs within vessels might be negligible in LSCI. The potential impact of radially diffusive motion of RBCs within vessels to findings of this study is also discussed in section 4.1.

The simple vascular geometry for simulation is made of parallel and equally spaced vessels (similar to ref. [39]). Vessels in the first geometry are parallel to *y* axis, and parallel to *z*-axis in the second geometry (Fig. 2a). Some of the major settings are as follows: geometry size: 1 × 1 × 1mm^3^, the radius of vessels: 0.01 mm, space between vessels: 0.1 mm, radius of the incident beam: 0.25 mm, radius of the detector: 0.01 mm, NA of the detector: 0.2. The detector is placed in the center of the field of view and right above the vessels. The optical properties of vessels and tissue are the same as in ref. [39].

For the realistic vascular geometry (shown in Fig. 2b), the simulation flow for generating the photon trajectories through the 3D geometry is detailed by Jafari et al.^28^. Briefly, parallelized Dynamic Light Scattering Monte Carlo (DLS-MC) simulations were launched on the Stampede2 Skylake compute nodes on Texas Advanced Computing Center (TACC) using the Message Passing Interface (MPI) protocol to simulate 80 × 10^9^ photon trajectories through the geometry.

The realistic vascular geometry was obtained through 2P imaging, followed by vectorization of the vascular structure^40^. The vectorized geometry was voxelized into a three-dimensional matrix of the size 277 × 277 × 303 voxels in the X, Y, and Z directions, respectively. The voxel size was a cubic 2 × 2 × 2 *μ*m^3^, yielding a total geometry size of 554 × 554 × 606 *μ*m^3^. A circular collimated wide-field beam with a flat profile was set to illuminate 95% of the top surface of the geometry. A NA of 0.25 and detector size of 9.8 × 9.8 *μ*m^2^ were used in the simulation settings to reflect the typical configuration of LSCI experimental setup. For detected photons, both entry and exit locations as well as the photon trajectories through the volume and photon weights were recorded. The optical properties of capillaries, non-capillary vessels and extra-vascular tissue as used in ref. [28] are adopted.

### 2.4. In vivo experimental validation

The MESI imaging system detailed in ref. [21, 22] is used for microfluidics and *in vivo* speckle imaging experiments. The laser wavelength is 785 nm and the magnification of the system is 2x. 56 MESI sequences with each sequence containing 15 speckle contrast images corresponding to 15 exposure times are acquired. The *K*^2^ curves are calculated by averaging speckle contrast values over multiple MESI sequences and then squaring it. ICT values are then extracted by fitting the *K*^2^ curves based on the MESI model^22^.

The mouse cranial window preparation procedures were detailed by Kazmi et al.^26^. During imaging sessions, the mouse (C57BL/6, Charles River Laboratories Inc.) was anesthetized with medical grade O_2_ vaporized isoflurane (3% induction, 1.5% maintenance).

For 2P imaging, images were acquired with a custom microscope and laser system^42,43^. The same anesthesia procedure as above-mentioned was used. In addition, 100 *μ*L of 70 kDa dextran-conjugated Texas Red diluted in saline at a 5% w/v ratio was added to the blood plasma through retro-orbital injection prior to imaging. The dye was then excited by an Yb fiber amplifier (λ = 1060 nm). 30 frames acquired with a resonant scanner were averaged at each depth to produce images.

All animal procedures in this study were approved by The University of Texas at Austin Institutional Animal Care and Use Committee (IACUC).

## 3. Results

### 3.1. Monte Carlo Simulation

The simulation results on the realistic vascular network under normal illumination are shown in Fig. 3. Figure 3a-e shows the map of the number of detected dynamic photons (i.e., photons experiencing dynamic scattering at least once), Var (*k*), E (*k*), and Var (*Y*), respectively. To counter the effects of wavelength, the Var (*Y*), Var (*k*) and E (*k*) are normalized by the wavenumber, i.e. Var(*Y/k*_0_), Var(*k/k*_0_) and E(*k/k*_0_), unless specified otherwise.

**Figure 3:**
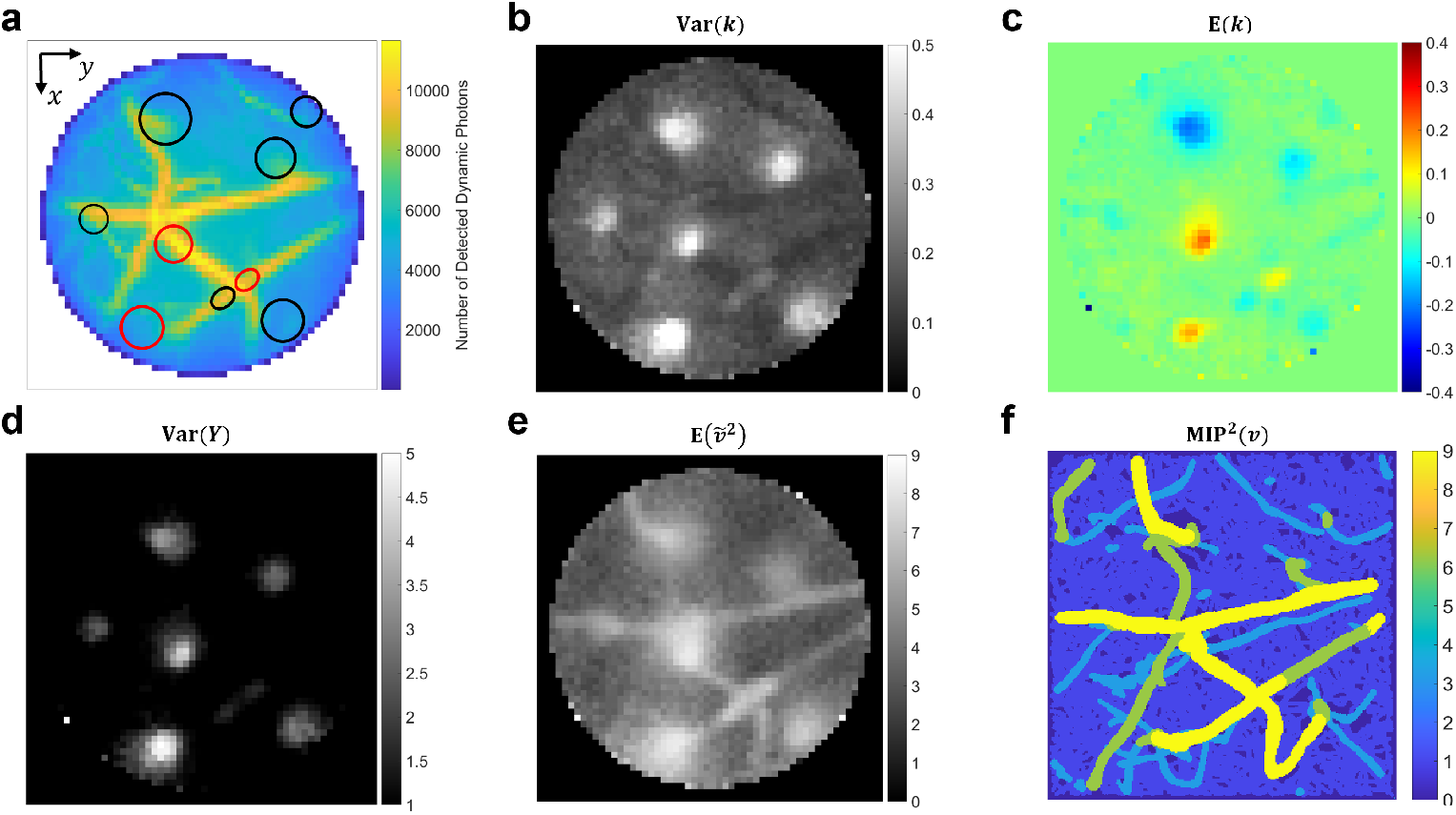
Simulation results on the realistic vascular network under normal illumination. **a** The map of the number of detected dynamic photons at each camera pixel. The dynamic photons refer to photons which experience at least one dynamic scattering event before exiting and being detected. **b-f** The map of Var(k), Mean (*k*), Var (*Y*), 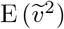, and MIP^2^ (*v*), respectively. MIP^2^ (*v*) represents the square of maximum intensity projection of flow speeds assigned in simulation. The position of bright spots in (**b**), (**c**) and (**d**) is circled out correspondingly in (**a**).

There are several interesting observations. First, note the bright blobs in Fig. 3b, d and blue/orange blobs in Fig. 3c whose positions are circled in Fig. 3a. The reason that those blobs appear in those places instead of elsewhere is suspected to be correlated with the underlying vascular structure and we find that there is always a major descending/ascending vessel in those spots (Fig. 2b and Supplemental Material section 5).

Second, the major surface vessels extending in the *x-y* plane shown in Fig. 3a appear even darker than surrounding parenchyma regions in the Var (*k*) map (Fig. 3b). This suggests that Var(*k*) might be sensitive to the orientation of vessels, namely, small for *x-y* plane surface vessels while large for *z*-directional descending vessels under normal illumination. This hypothesis is verified by simulation results on the simple vascular geometries consisting of parallel and equally-spaced vessels. Var (*k*) of *z*-directional vessels is observed to increase by 7 times compared with that of *y*-directional vessels under normal illumination (Supplemental Material section 6).

Third, among those blobs, some are blue while others appear orange in Fig. 3c. Further analysis on the flow vector assignment in simulation reveals that this is dependent on whether the flow vector is z-positive or z-negative (Supplemental Material section 7).

Finally, E (*k*) of parenchyma regions is 0 (Fig. 3c). Hence, Eq. 9 can be applied with good accuracy. With Var (*Y_s_*) being 0 in simulation, we could calculate 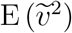 by dividing Var (*Y*) with Var (*k*). As Fig. 3e shows, the 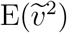 map reveals a clearer structure of surface vessels than Var (*Y*) (Fig. 3d). To evaluate accuracy of the absolute blood flow speed estimation given in Fig. 3e, the square of the maximum intensity projection (MIP) of blood flow speeds assigned in simulation is mapped in Fig. 3f. In terms of the resolved vessels, the shape of *x-y* plane surface vessels is well preserved in Fig. 3e and the intensity value also shows a good match with that in Fig. 3f. However, for *z*-directional descending vessels, the vessel boundary is expanded and there is no clear border, which highlights their distinct properties in LSCI from *x-y* plane surface vessels.

### 3.2. Experimental validation

The enhanced sensitivity of speckle contrast to blood flow speed in *z*-directional vessels compared with *x-y* plane vessels is also observed *in vivo*. As highlighted by white arrows in Fig. 4a and b, the orthogonal X-Y and X-Z cross-sections of 2P imaging data clearly reveal an inverted “L” shaped vessel. At the end of its surface strand, it develops into a descending strand into the tissue. ROI of the two strands is shown by the white boxes in Fig. 4d. Notice that there is no other major vessel branch on this “L” shape and vessel diameters of the two strands are approximately the same. Thus, based on blood flow conservation^26^, the flow speed should be approximately the same in these two strands. Nevertheless, we see a lower speckle contrast in areas corresponding to the descending strand as pointed out by the white arrow in Fig. 4c. It indicates an enhancement of the sensitivity of speckle contrast to blood flow speed in the descending strand compared with surface strand. More specifically, the average ICT squared of the descending strand is ~78% larger than that of its surface counterpart, as highlighted by white boxes in Fig. 4d). If the impact of Var(*Y_s_*) is negligible, it implies that Var (*k*) of the descending strand would be more than 75% larger than that of surface strand. Notably, similar border expansion is also observed here in the position of the descending strand as in simulation (Fig. 3e).

**Figure 4:**
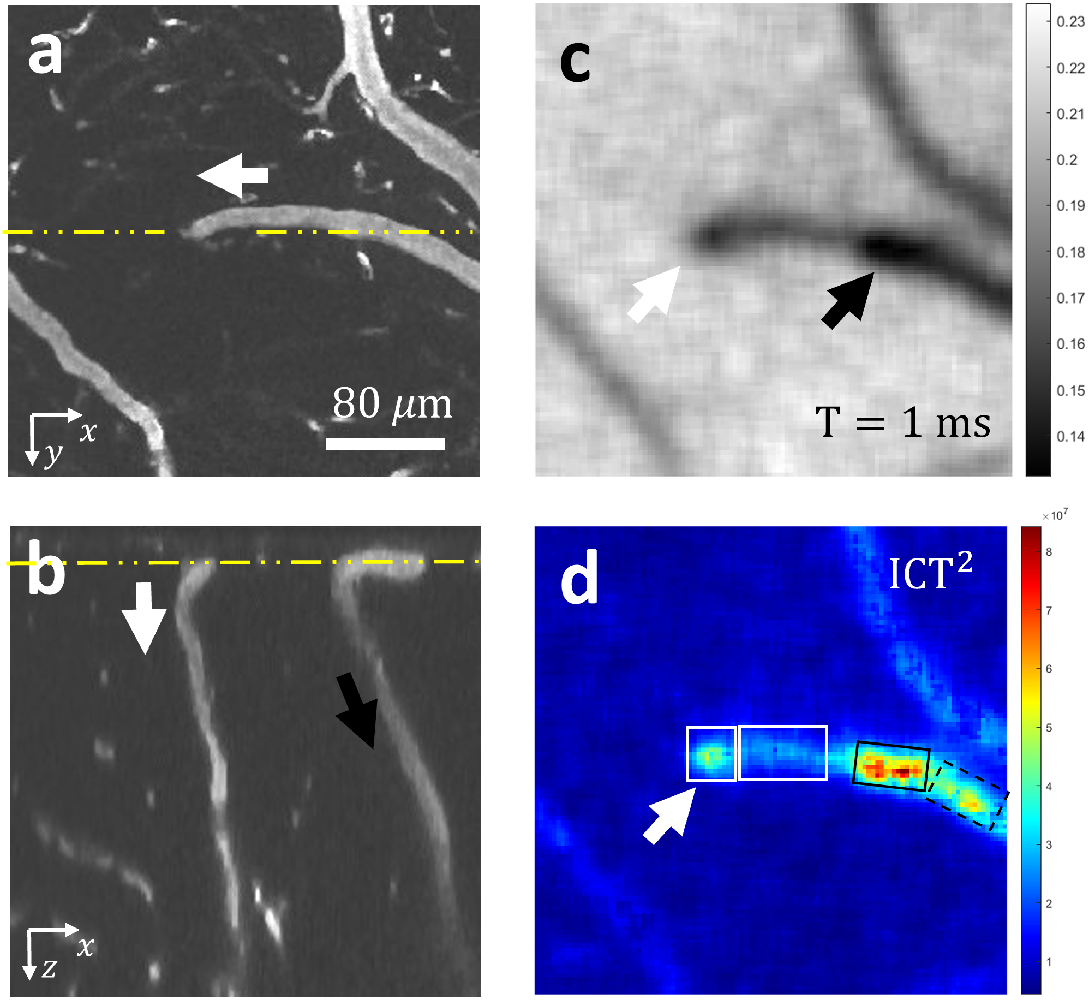
One example of the *z*-directional descending vessel inducing a more significant decrease of speckle contrast and increase of ICT than its upstream *x-y* plane surface strand *in vivo*. **a**, **b** Orthogonal cross-sections of the vascular structure acquired from 2P imaging. (**a**) X-Y cross-section (depth: 27 *μ*m); (**b**) X-Z cross-section along the yellow dashed line in (**a**). The yellow line in (**b**) indicates the *z* position of (**a**). White arrows in (**a**) and (**b**) indicate the direction of an upside-down “L” shaped vessel which has a surface strand extending horizontally on the surface followed by a descending strand deep into the tissue. scale bar: 80 *μ*m. The spatial scale along the x- and z-axis is the same in (**b**). **c** Single-exposure speckle contrast image of the vascular network. Camera exposure time T=1 ms. The white arrow indicates position of the descending vessel strand which shows a stronger decrease of speckle contrast than its connected *x-y* plane surface strand. **d** The map of ICT square. The two white boxes highlight the descending strand and surface strand, respectively. The white arrow indicates the position of the descending strand exhibiting a larger ICT value than its connected *x-y* plane counterpart. The black arrow in **b** highlights another descending vessel branch from the main vessel. The black arrow in **c** and black boxes in **d** indicate the compound effects of vascular structure and blood flow in modulating speckle contrast and ICT values.

To further evaluate the statistical significance of the enhanced sensitivity, 9 pairs of *z*-directional vessels and *x-y* plane vessels from 3 mice are analyzed. Those vessel pairs are selected for analysis because their vascular structure has the same properties as in the example mentioned above, i.e., the upside-down L vessel shape and the approximately same diameter of the two strands. The location of four *z* and *x-y* strand pairs in a typical mouse cerebral window is shown in Fig. 5a and their vascular structure is acquired by 2P imaging (Fig. 5b). ROI of the *z* and *x-y* strands in each pair is selected in the similar way as shown by white boxes in Fig. 4d. The ICT squared of the *z* strand in each pair is plotted against that of the *x-y* strand (Fig. 5c). The linear fitting results in a slope of approximately 2. In addition, the difference between ICT squared of *z* strand and that of its contiguous *x-y* strand is statistically significant (paired t-test, n=9, *p* < 0.001). Those results provide direct experimental evidence for the enhanced sensitivity of ICT to blood flow changes in *z*-directional vessels compared with vessels extending in the *x-y* plane.

**Figure 5:**
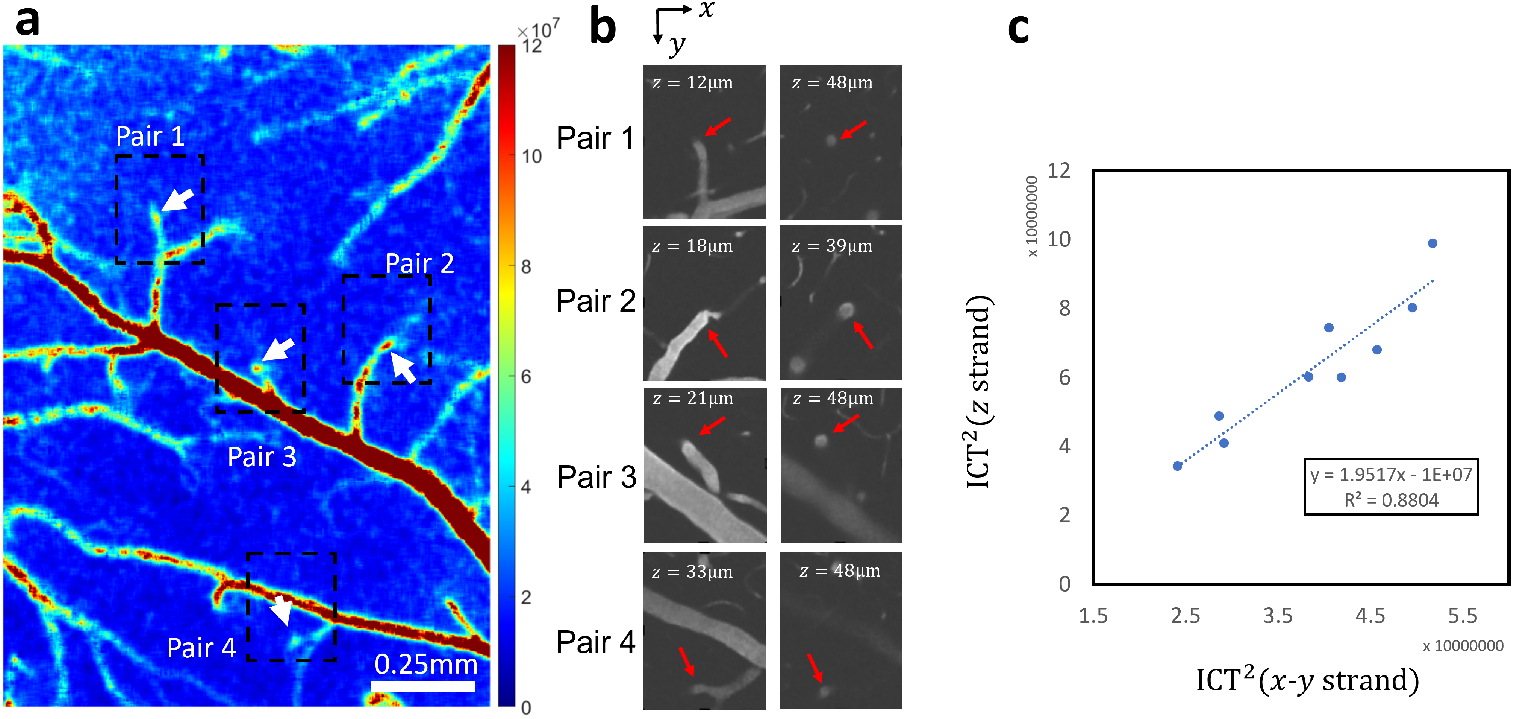
Statistical comparison of ICTs between the *z*-directional descending vessel strand and its upstream *x-y* plane surface strand *in vivo*. **a** Pairs of *z* strand and *x-y* strand in a typical mouse cerebral window whose position is highlighted by the dashed rectangle on the ICT squared image. The arrows point to the descending vessel in each pair. Scale bar: 0.25mm. **b** The corresponding 2P images showing the vascular structure of the four pairs of *z* strands and *x-y* strands. The first column shows the X-Y cross-section of the vascular structure near the surface while the second column shows the cross-section at the same location but in a deeper tissue. The red arrows point to the *z* strands in the pair. **c** The juxtaposing of ICT squared of the *z* strand and that of the contiguous *x-y* strand. The scatter plot shows the data of nine *z* and *x-y* strand pairs from three mice. The linear fitting is performed and the slope is around 2, larger than 1, which indicates that the ICT of *z*-directional vessels could be larger than that of *x-y* plane vessels even though they assume the same blood flow speed. The difference between ICT squared of the *z* strand and that of the paired *x-y* strand is statistically significant (paired t-test, n=9, *p* < 0.001).

Finally, the compound effects of vascular structure and blood flow should be noticed. As highlighted by the black arrow in Fig. 4b, there is another descending vessel branch from the main vessel. The blood flow in the main vessel splits into two portions: one goes into the above-mentioned inverted “L” shaped vessel and the other goes into this second descending vessel. Therefore, the speckle contrast and ICT in the solid black rectangle in Fig. 4d result from not only an underlying descending vessel but also the larger blood flow in the main vessel. That is why they are not directly comparable with those of the descending strand in the white square in Fig. 4d. Interestingly, the enhanced sensitivity can be roughly examined if we compare the ICT squared in the solid-line black rectangle with that in the dashed black rectangle in Fig. 4d. Both areas cover the main vessel of the largest blood flow but the solid rectangle assumes larger ICT squared, which indicates the additional impact of the descending vessel in the solid square. Similarly, the joint effects of vascular structure and blood flow are observed in the region of vessel pair 2 in Fig. 5a where a second descending vessel is also present (Fig. 5b).

## 4. Discussion

### 4.1. Bridging ICT to physiologically meaningful blood flow variables

Given the volume integrated nature of LSCI and the complexity of the vascular network, it has been long hypothesized that if LSCI is measuring some physiologically meaningful blood flow variable, it measures the weighted average of that variable within the probed volume [22, 39]. However, it is unclear how the weighting is determined. Our derivation in section 3.1 and 3.2 reveals that the weighting is determined by photons’ dynamic scattering process, according to the definition of 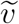 (Eq. 6). Note that 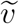 is physiologically meaningful and represents the weighted average of blood flow speeds probed by detected photons. In the case that all dynamic scattering events sample the same blood flow speed *v*, 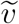 would be equal to *v* regardless the weighting.

Combining the results in section 2.1 and 2.2, a general relationship between ICT and physiologically meaningful blood flow variables, i.e. 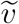, can be established. In section 2.1, it has been shown that ICT squared is pro-portional to the variance of *Y*, which accommodates all current *g*_1_(*τ*) models in LSCI. In section 2.2, Eq. 9 further points out that the variance of *Y* assumes a linear relationship with the expectation of 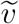 squared. Hence, the linear relationship between ICT squared and the expectation of 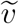 squared is reached by our theoretical derivation.

Contrary to the frequently adopted notion that ICT is proportional to blood flow speed, our theory concludes that the squares of the two that are linear to each other. The difference between the two notions is centered on whether Var (*Y_s_*) is zero or not. Pragmatically, the existence of Var (*Y_s_*) is indispensable to tackling the “biological zero” problem which refers to the non-zero residual signal even when no blood flow is present^37,44^. A non-zero Var (*Y_s_*) might reduce the difference of ICT values between descending/ascending vessels and surface vessels. As observed in simulation, the Var (*k*) of *z*-directional vessels is 7 times larger than that of *y*-directional vessels in the simple vascular geometry, which is expected to generate a ICT squared difference of 7 times according to Eq. 9 if Var (*Y_s_*) is zero. However, ICT squared *in vivo* is only 60% larger on average in descending/ascending vessels than vessels laying on the surface. A non-zero Var (*Y_s_*) *in vivo* could play a significant role in accounting for such discrepancy. In addition, the scatters’ movement could be more diverse *in vivo* compared with our settings in simulation. For example, the radially diffusive motion of RBCs in vessels is not present in our simulation.

Var (*k*) plays a major role in bridging ICT and the physiologically meaningful blood flow variable, 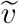, as Eq. 9 points out. It provides a theoretical basis for extracting the physiologically meaningful blood flow variable 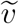 from ICT. Given the Var (*k*) already known of the vascular network, the estimation of absolute statistical blood flow speeds could be made by dividing the measured Var (*Y*) by Var (*k*) (Fig. 3e). Note that Var (*Y_s_*) is assumed 0 in simulation.

Finally, our findings showed partial support for Briers et al.’s method of converting measured ICT values to absolute blood flow speeds. Specifically, the conversion is performed by *v_c_* = λ/(2*πτ_c_*) where *v_c_* is named decorrelation velocity, λ is wavelength and *τ_c_* is correlation time^16,17^. Our theory shows that this is in fact acquired by assuming Var (*Y_s_*) = 0 and approximating Var (*k*) with 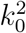, the square of wavenumber (Eq. 9).

### 4.2. *Interpretation and practical implications of* Var (*k*)

Var (*k*) can be interpreted at both the microscopic and macroscopic scales. Microscopically, by definition (Eq. 6 and 8), it provides an essential characterization for the variation of photons’ dynamic scattering process inside the probed medium in terms of the accumulated frequency shift. Macro-scopically, as revealed by Eq. 9, it reflects the ability of the probed vascular network to induce a decrease of speckle contrast or increase of ICT for a given illumination and detection setup. The micro and macro-scale interpretations are connected due to the fact that photons’ dynamic scattering is mainly constrained within the vascular network.

Var (*k*) also illustrates that it is challenging, if not impossible, to do absolute blood flow speed measurements through a generalized calibration since Var (*k*) is unique for a given vascular network. When the probed vasculature changes, Var (*k*) also changes.

### 4.3. Physical mechanism and practical implications of the directionality susceptibility

The different sensitivity of ICT to the blood flow speed in *x-y* plane vessels and *z*-directional descending/ascending vessels is likely due to the different flow direction in those vessels. One might argue that in descending/ascending vessels, the ratio of blood volume in the overall volume sampled by detected photons might be larger than that in the surface vessels. And it might be the larger blood volume ratio in descending/ascending vessels that is causing the higher sensitivity of ICT to the blood flow. We exclude this theory by manipulating the flow direction in simulation. If the blood volume ratio theory is true, then Var (*k*) of *z*-directional vessels should remain larger than that of *x-y* plane vessels even if the direction of flow is changed since the blood volume ratio is invariant. However, it is observed that Var (*k*) of the *z*-directional vascular geometry is smaller than that of *y*-direction vascular geometry after switching the flow direction in *z*-directional vascular geometry to *y*-direction and that in *y*-directional vascular geometry to *z*-direction (Table S2). Validation on the realistic vascular geometry shows consistent results (Fig. 6). The bright blobs originally present in the Var (*k*) map (Fig. 3b) are removed after the amplitudes of *y* and *z* component of the unit velocity vector of blood flow are swapped (Fig. 6a). Instead, the position of *y*-directional vessels is highlighted. However, such phenomena would not appear if it is the amplitudes of *x* and *y* component that are swapped (Fig. 6b).

**Figure 6:**
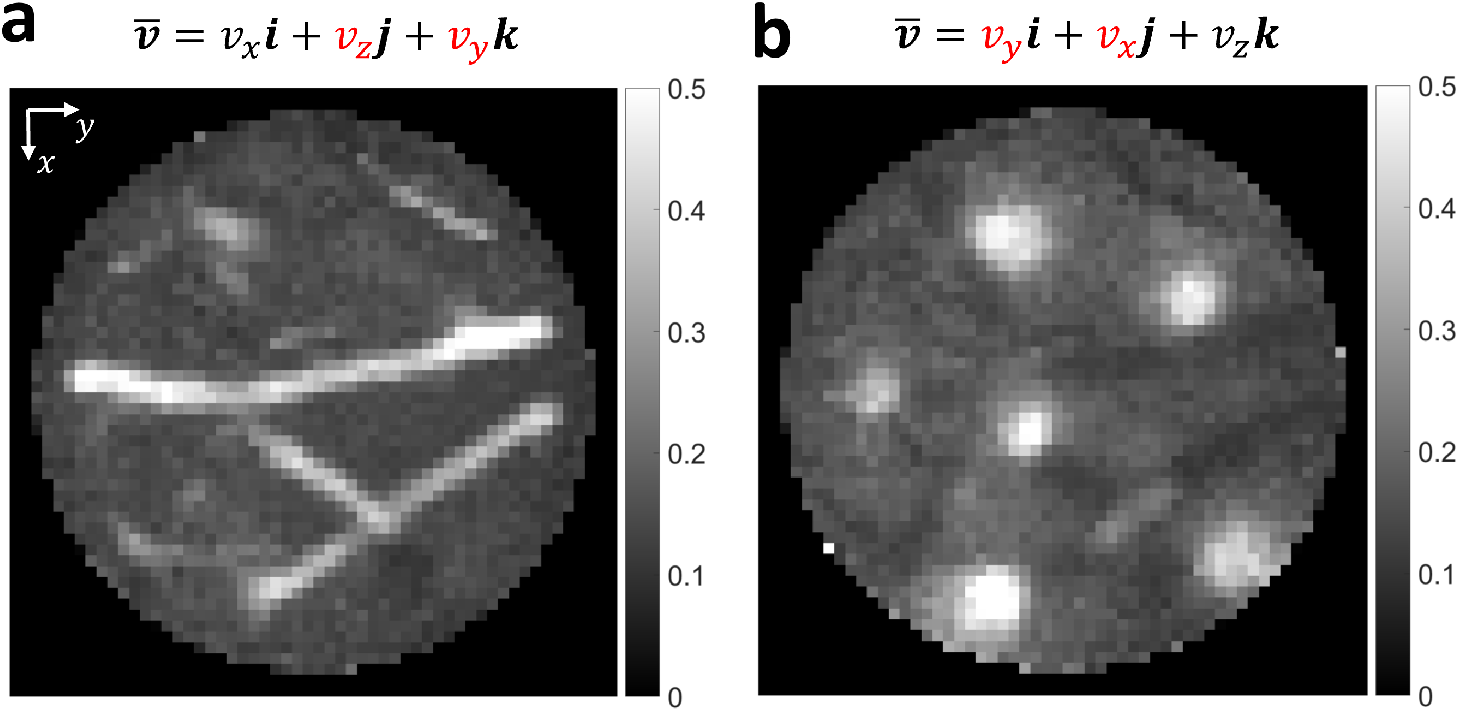
Map of Var (*k*) with the blood flow direction switched in realistic vascular geometry. **a** The amplitudes of *y* and *z* components of the unit velocity vector of blood flow are swapped in simulation. **b** The amplitudes of *x* and *y* components of the unit velocity vector of blood flow are swapped in simulation.

Var (*k*) not only quantifies and reveals the directionality susceptibility of LSCI to vessel orientation, but also helps explain such susceptibility. Since Var (*k*) measures the overall variance of *k* of detected photons, it is susceptible to the range determined by the extreme values of *k*. Consider extreme values of *k* of singly scattered photons first where 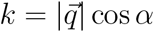. Under normal illumination, 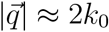 and 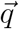 is along the *z*-axis. Therefore, *k* would be maximized when cos *α* is maximized. Since *α* is the angle between velocity vector 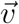 and momentum transfer vector 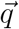 which is along the *z*-axis, cos *α* would be maximized when 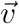 is also along the *z*-axis. That explains why *z*-directional vessels have larger Var (*k*) values than *x-y* plane surface vessels. Finally, for photons that are scattered multiple times, the randomized directions of 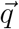 and 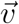 make it difficult to generate extreme *k* values since the effects of multiple scattering events can cancel out with each other. Hence, the directionality susceptibility mainly originates from the directionality susceptibility retained in single-scattering or few-scattering components of the detected light.

The directionality susceptibility of LSCI has several practical implications. First, direct comparison of ICT values in *x-y* plane vessels and in ascending/descending vessels should be avoided. If a descending/ascending vessel exhibits a larger ICT value, it does not necessarily imply a higher blood flow than its surface counterpart. Second, the special property of *z*-directional vessels, i.e., enhanced ability to induce the decrease of speckle contrast and increase of ICT under the same blood flow speed, might be useful in locating descending/ascending vessels. Note that those types of vessels are particularly prevalent in cerebral cortex and play an important role in the blood supply to deeper tissues.

## 5. Conclusion

The interpretation of ICT is a key topic in quantitative LSCI. Though it has been mainly considered as a metric of blood flow, there is increasing evidence that it is susceptible to the structure of vascular network. We build a theoretical framework to facilitate the modeling of the vascular network’s impact on ICT and find that ICT is modulated by vessel orientation. In both simulation and *in vivo* experimental validation, ICT of descending/ascending vessels exhibits a higher sensitivity to blood flow changes than in surface-extending vessels. The different sensitivity is shown due to the flow direction instead of blood volume ratio by simulation. The single-scattering component of the detected light might play a major role in ICT’s susceptibility to vessel orientation. Those results suggest that the impact of vascular network structure warrants more attention and investigation in the interpretation of ICT.

## Acknowledgements

This work is supported by National Institutes of Health (NIH) (Grant NS108484, EB011556) and UT Austin Portugal Program. The authors acknowledge Texas Advanced Computing Center (TACC) at UT Austin for providing high-performance computing resources. The authors also would like to express thanks to Chenmu Zhang and Dr. Zhongcan Xiao for the inspiring discussion on Supplemental Material section 3; thanks to Dr. Colin Sullender for the advice on troubleshooting MESI system; and finally, thanks to Samuel A. Mihelic for the help with Blender rendering.

## Author Contributions

Q.F. and A.K.D. developed the theory and designed the experiments; Q.F. and C.Z.J did the Monte Carlo simulation experiments; S.E. and A.T performed the *in vivo* mice imaging experiments; Q.F., C.Z.J, A.T., S.E. and A.K.D wrote the paper together.

