## Supplemental Material for "Interpreting Inverse Correlation Time: from Blood flow to Vascular Network"

#### Section 1. Why can $Y$ be viewed as the accumulation of frequency shifts in dynamic scattering?

Proof:

Set up the coordinate system as the diagram on the right shows: the photon starts from point S, bumps into the scattering particle at point B and finally arrives at point P. The velocity vector of the scattering particle is $\boldsymbol{v}$ and the angle between $\overrightarrow{SB}$ and $\boldsymbol{v}$ is α while β between $\boldsymbol{v}$ and $\overrightarrow{BP}$. Assume that the wavenumber of the incident light is $k_0$, the light speed is $c$, the original frequency of the light is $f$, and the momentum transfer vector is defined as $\boldsymbol{q} = \boldsymbol{k}_o - \boldsymbol{k}_i$ as in the article.

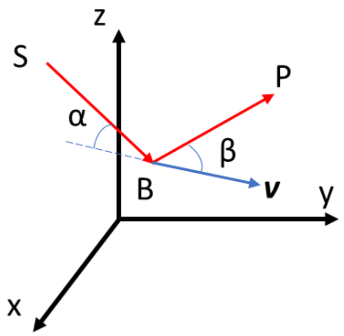


According to Doppler effect, the frequency observed on the scattering particle

$$f_1 = \frac{c - |\boldsymbol{v}|cos\alpha}{c} f$$

Since $cos\alpha = \frac{\overrightarrow{SB}\cdot\boldsymbol{v}}{|\overrightarrow{SB}||\boldsymbol{v}|}$,

$$f_1 = \frac{c - \frac{\overrightarrow{SB}\cdot\boldsymbol{v}}{|\overrightarrow{SB}|}}{c} f$$

Similarly, the frequency of the scattered light observed at point P is

$$f_2 = \frac{c}{c - \frac{\overrightarrow{BP}\cdot\boldsymbol{v}}{|\overrightarrow{BP}|}} f_1$$

Thus, we have

$$f_2 = \frac{c - \frac{\overrightarrow{SB}\cdot\boldsymbol{v}}{|\overrightarrow{SB}|}}{c - \frac{\overrightarrow{BP}\cdot\boldsymbol{v}}{|\overrightarrow{BP}|}} f = (1 + \frac{\frac{\overrightarrow{BP}\cdot\boldsymbol{v}}{|\overrightarrow{BP}|} - \frac{\overrightarrow{SB}\cdot\boldsymbol{v}}{|\overrightarrow{SB}|}}{c - \frac{\overrightarrow{BP}\cdot\boldsymbol{v}}{|\overrightarrow{BP}|}}) f$$

Since the speed of the scattering particle is much smaller than that of light in blood flow imaging scenarios, we have $\frac{\overrightarrow{BP}\cdot\boldsymbol{v}}{|\overrightarrow{BP}|} \approx 0$ to arrive at the following approximation

$$f_2 = (1 + \frac{\frac{\overrightarrow{BP}\cdot\boldsymbol{v}}{|\overrightarrow{BP}|} - \frac{\overrightarrow{SB}\cdot\boldsymbol{v}}{|\overrightarrow{SB}|}}{c}) f$$

Therefore, the frequency shift is

$$\Delta f = f_2 - f = \frac{(\frac{\overrightarrow{BP}}{|\overrightarrow{BP}|} - \frac{\overrightarrow{SB}}{|\overrightarrow{SB}|}) \cdot \boldsymbol{v}}{c} f$$

Given that

$$\boldsymbol{q} = \boldsymbol{k}_o - \boldsymbol{k}_i = k_0(\frac{\overrightarrow{BP}}{|\overrightarrow{BP}|} - \frac{\overrightarrow{SB}}{|\overrightarrow{SB}|})$$

We can rewrite $\Delta f$ as

$$\Delta f = \frac{\boldsymbol{q} \cdot \boldsymbol{v}}{k_0 c} f = \frac{\boldsymbol{q} \cdot \boldsymbol{v}}{2\pi f} f = \frac{\boldsymbol{q} \cdot \boldsymbol{v}}{2\pi}$$

Hence

$$\Delta\omega = \boldsymbol{q} \cdot \boldsymbol{v}$$

The conclusion is the same for the case where the particle velocity vector $\boldsymbol{v}$ is on the inverse direction of $\overrightarrow{SB}$, namely, the two is heading towards each other.

Considering that $Y = \sum_{i=1}^{N} \boldsymbol{q}_i \boldsymbol{V}_i$ in definition, $Y$ can be viewed as the accumulation of frequency shifts in dynamic scattering.

**Section 2. Relating $\boldsymbol{\sigma}$ to $\frac{1}{\tau_c}$ for the Gaussian $\boldsymbol{g_1}$ model using derivatives of Fourier transform**

Assuming the Fourier transform of $g_1(\tau)$ is $P(Y)$, namely,

$$g_1(\tau) = e^{-(\tau/\tau_c)^2} \xrightarrow{F.T.} P(Y)$$

Then the Fourier transform of the derivative of $g_1(\tau)$ is

$$g_1{}'(\tau) \xrightarrow{F.T.} jYP(Y)$$

Similarly, the Fourier transform of the second-order derivative of $g_1(\tau)$ is

$$g_1{}''(\tau) \xrightarrow{F.T.} -Y^2 P(Y)$$

Note that $\sigma$ is the square root of the variance of $P(Y)$ while the variance of $P(Y)$ is given by

$$Var(Y) = \int_{-\infty}^{\infty} Y^2 P(Y)\, dY$$

Therefore,

$$\sigma^2 = Var(Y) = -g_1{}''(\tau = 0)$$

Given that

$$g_1''(\tau) = e^{-\left(\frac{\tau}{\tau_c}\right)^2} (\frac{4\tau^2}{\tau_c{}^4} - \frac{2}{\tau_c{}^2})$$

We arrive that $\sigma^2 = -g_1''(\tau = 0) = \frac{2}{{\tau_c}^2}$

Therefore,

$$\frac{1}{\tau_c} = \frac{1}{\sqrt{2}}\sigma$$

**Section 3. Why does $Var(Y) \propto (\frac{1}{\tau_c})^2$ hold for any electric field auto-correlation functions taking the form of $g_1(\tau) = e^{-(|\tau|/\tau_c)^n}$ as long as it is agreed that the spectrum of detected light is both bandwidth- and amplitude-limited in reality?**

Proof:

Assume the Fourier transform pair

$$g(x) = e^{-(|x|)^n} \xrightarrow{F.T.} P(Y)$$

Then according to the scaling property of Fourier transform, assuming $\tau_c > 0$ we have

$$g(x/\tau_c) = e^{-(|x/\tau_c|)^n} \xrightarrow{F.T.} \tau_c P(\tau_c Y)$$

Assuming the spectrum of detected light is both bandwidth- and amplitude-limited, that is

$$P(Y) = 0, \forall |Y| > B/2$$

$$P(Y) < C, \forall |Y| \le B/2$$

then we have

$$Var_0(Y) = \int_{-\infty}^{\infty} Y^2 P(Y) dY < \int_{-B/2}^{B/2} Y^2 C dY = \frac{B^3 C}{12}$$

Namely, $Var_0(Y)$ is finite.

Therefore, the variance of the scaled spectrum is

$$Var(Y) = \int_{-\infty}^{\infty} Y^2 \tau_c P(\tau_c Y) dY$$

Applying substitution $u = \tau_c Y$, we have

$$Var(Y) = \int_{-\infty}^{\infty} \frac{1}{{\tau_c}^2} u^2 \tau_c P(u) \frac{1}{\tau_c} du = \frac{1}{{\tau_c}^2} \int_{-\infty}^{\infty} u^2 \, P(u) du = \frac{1}{{\tau_c}^2} Var_0(Y)$$

Therefore, $Var(Y) \propto \frac{1}{{\tau_c}^2}$. The proof is over.

Brief discussion:

We should notice that the constant $Var_0(Y)$ is dependent on the shape of the spectrum, namely, determined by the modulation number $n$ in $g_1(\tau) = e^{-(|\tau|/\tau_c)^n}$. We also highlight that the advantage of interpreting ICT in the frequency domain is that it allows decoupling effects of ICT and modulation number $n$ into the product of each one's, given $Var(Y)$ is just the product of $\frac{1}{{\tau_c}^2}$ and $Var_0(Y)$ while in time domain, the effects of ICT and $n$ on $g_1$ are coupled non-linearly.

**Section 4. Why would $\tilde{v}$ and $k$ be uncorrelated given that the blood flow speeds are uncorrelated with the photon's scattering geometry?**

By definition,

$$k = \sum_{i=1}^{N} |\boldsymbol{q}_i| cos\alpha_i$$

$$\tilde{v} = \frac{\sum_{i=1}^{N} |\boldsymbol{q}_i| cos\alpha_i \cdot |\boldsymbol{v}_i|}{\sum_{i=1}^{N} |\boldsymbol{q}_i| cos\alpha_i}$$

Let $w_i = |\boldsymbol{q}_i| cos\alpha_i = 2k_0 \sin(\theta_i/2) cos\alpha_i$ where $\theta_i$ is the angle between the incident wavevector and scattered wavevector in the i-th dynamic scattering event, $v_i = |\boldsymbol{v}_i|$, then

$$k = \sum_{i=1}^{N} w_i$$

$$\tilde{v} = \frac{\sum_{i=1}^{N} w_i \cdot v_i}{\sum_{i=1}^{N} w_i}$$

If the blood flow speeds are uncorrelated with the photon's scattering geometry, i.e., $|\boldsymbol{v}|$ is uncorrelated with $\theta$ and $\alpha$, then $w$ and $v$ would be uncorrelated, which means

$$E(wv) = E(w)E(v)$$

Let $w_1, w_2, \dots, w_N$ be N independent and identically distributed (i.i.d.) random variables, and their expectation is $E(w)$. Similarly, let $v_1, v_2, \dots, v_N$ be N i.i.d random variables and their expectation is denoted as $E(v)$. In addition, $w$ and $v$ is uncorrelated.

Then what we need to prove is that $E(k\tilde{v}) = E(k)E(\tilde{v})$.

Proof:

Since

$$k = \sum_{i=1}^{N} w_i$$

$$\tilde{v} = \frac{\sum_{i=1}^{N} w_i \cdot v_i}{\sum_{i=1}^{N} w_i}$$

We have

$$E(k\tilde{v}) = E\left(\sum_{i=1}^{N} w_i \cdot v_i\right) = \sum_{i=1}^{N} E(w_i \cdot v_i) = \sum_{i=1}^{N} E(w_i)E(v_i) = \sum_{i=1}^{N} E(w)E(v) = NE(w)E(v)$$

$$E(k) = E\left(\sum_{i=1}^{N} w_i\right) = \sum_{i=1}^{N} E(w_i) = \sum_{i=1}^{N} E(w) = NE(w)$$

$$E(\tilde{v}) = E\left(\frac{\sum_{i=1}^{N} w_i \cdot v_i}{\sum_{i=1}^{N} w_i}\right) = E\left(\sum_{i=1}^{N} \frac{w_i}{\sum_{i=1}^{N} w_i} \cdot v_i\right)$$
$$= \sum_{i=1}^{N} E\left(\frac{w_i}{\sum_{i=1}^{N} w_i} \cdot v_i\right) = \sum_{i=1}^{N} E\left(\frac{w_i}{\sum_{i=1}^{N} w_i}\right) E(v_i) = E(v) \sum_{i=1}^{N} E\left(\frac{w_i}{\sum_{i=1}^{N} w_i}\right)$$
$$= E(v)E\left(\sum_{i=1}^{N} \frac{w_i}{\sum_{i=1}^{N} w_i}\right) = E(v)E\left(\frac{\sum_{i=1}^{N} w_i}{\sum_{i=1}^{N} w_i}\right) = E(v)$$

Therefore, $E(k\tilde{v}) = E(k)E(\tilde{v})$, namely, $k$ and $\tilde{v}$ are uncorrelated.

**Section 5. The position of descending vessels in the realistic vascular geometry**

The major surface vessels (extending in the X-Y plane) are located close to the top surface of the geometry (**Fig. S1 A**) while the descending vessels (extending in the Z direction) dip from the top surface into the deeper tissue (Fig. **S1 B-D**). Comparing the positions of those descending vessels (highlighted by circles in **Fig. S1 C**) with those bright spots observed in **Fig. 3 b-d**, we can see that they show high coincidence.

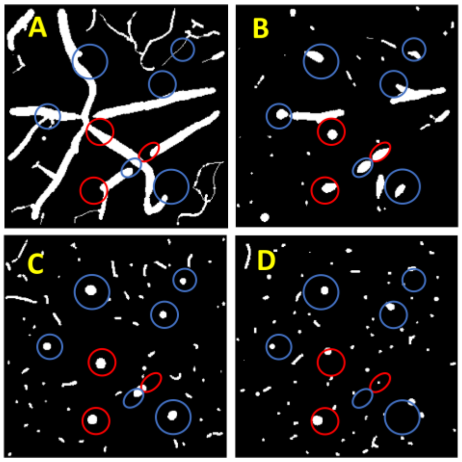


**Figure S1. The position of descending vessels in the realistic vascular geometry.** A-D show the X-Y cross-sections of the vascular geometry matrix in the Z layer of 21, 29, 37, 102, respectively. The distance between two consecutive Z layers is 3 μm. The overall size of the vascular geometry matrix is 277×277×303.

**Section 6. $Var(k)$ of z-directional vessels is increased 7 times than that of y-directional vessels under normal illumination**

As **Table S1** highlights, the $Var(k)$ of the z-directional vessel geometry is 8 times that of the y-directional vessel geometry when the detector is placed right above the vessel. In addition, when the detector is placed above the parenchyma region, there is no significant difference between y- and z-directional vessel geometry in terms of $Var(k)$.

**Table S1. The simulation results on the simple vascular geometry using normal illumination.**

| Size: 3×3×3 mm^3<br>Resolution<br>ds=0.002mm<br>1e9 photons<br>launched | R(source)=0.25mm, $NA_s$=-0.25*, R(detector)=0.01mm, $NA_d$=0.2;<br>R(vessel)=0.01, $h_{space}$=0.1 | | | |
|---|---|---|---|---|
| | Detector position (-0.05, -0.05, 0)<br>Right above vessel | | Detector position (0, 0, 0)<br>Right above parenchyma | |
| | Y-directional vessel geom | Z-directional vessel geom | Y-directional vessel geom | Z-directional vessel geom |
| Mean(k/k0) | 0.0028 | -0.174 | 0.0001 | -0.0189 |
| Var(k/k0) | **0.0307** | **0.2516** | **0.0248** | **0.0225** |
| # of detected photons | 1131 | 1021 | 985 | 1093 |

*Note: a negative NA means collimated beam and the absolute value specifies the radius of the beam.

**Table S2. The simulation results on the simple vascular geometry using normal illumination.**

| Size: 3×3×3 mm^3<br>Resolution<br>ds=0.002mm<br>1e9 photons<br>launched | R(source)=0.25mm, $NA_s$=-0.25*, R(detector)=0.01mm, $NA_d$=0.2;<br>R(vessel)=0.01, $h_{space}$=0.1 | | | |
|---|---|---|---|---|
| | y-directional vessel geom (y-directional flow) | z-directional vessel geom (z-directional flow) | y-directional vessel geom (z-directional flow) | z-directional vessel geom (y-directional flow) |
| Mean(k/k0) | 0.0028 | -0.174 | -0.2260 | 0.0039 |
| Var(k/k0) | **0.0307** | **0.2516** | **0.3564** | **0.0126** |
| # of detected photons | 1131 | 1021 | 1131 | 1021 |

*Note: a negative NA means collimated beam and the absolute value specifies the radius of the beam.

### Section 7. The different behaviors of z-positive and z-negative flow vectors in terms of $E(\boldsymbol{k})$

**Fig. S3 A** highlights two adjacent descending vessel segments but whose signs of $E(k)$ are opposite as **Fig. 2C** shows. **Fig. S3 A** shows that their z-direction flow is different in that one is z-positive while the other is z-negative. Further analysis on the flow direction of

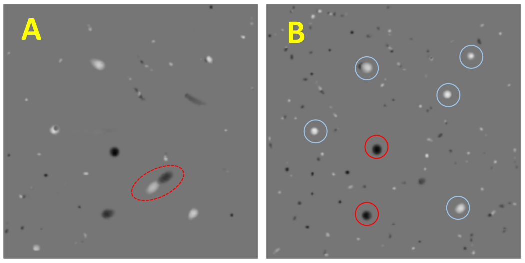


**Figure S3. The z-flow direction assignment of descending vessels.** Dark: z-negative, i.e., towards the top surface; bright: z-positive, i.e., away from the top surface. A and B correspond to z layer 31 and 40, respectively.

other descending vessels reveals the same principle that when the flow direction is z-positive, namely, the flow is away from the top surface, $E(k)$ is negative while when the flow direction is z-negative, i.e., the flow is towards the top surface, $E(k)$ is positive (**Fig. S3 B**).
